# Lymphoma cells are circulating in blood of Enzootic Bovine Leukosis and clonality value of virus-infected cells is a useful information for the diagnostic test

**DOI:** 10.1101/2022.06.07.495053

**Authors:** Md Belal Hossain, Tomoko Kobayashi, Sakurako Makimoto, Misaki Matsuo, Kohei Nishikaku, Benjy Jek Yang Tan, Akhinur Rahman, Samiul Alam Rajib, Kenji Sugata, Nagaki Ohnuki, Masumichi Saito, Toshiaki Inenaga, Kazuhiko Imakawa, Yorifumi Satou

**Affiliations:** Division of Genomics and Transcriptomics, Joint Research Center for Human Retrovirus Infection, Kumamoto University, Kumamoto, 860-8556, Japan; Department of Food Microbiology, Faculty of Nutrition and Food Science, Patuakhali Science and Technology University, Dumki, Patuakhali-8602, Bangladesh; Laboratory of Animal Health, Department of Animal Science, Faculty of Agriculture, Tokyo University of Agriculture, Atsugi, Kanagawa 243-0034, Japan; Department of Virology II, National Institute of Infectious Diseases, Tokyo, 162-8640, Japan.; Center for Emergency Preparedness and Response, National Institute of Infectious Diseases, Tokyo, 162-8640, Japan; Laboratory of Molecular Reproduction, Research Institute of Agriculture, Tokai University, Kumamoto 862-8652, Japan

## Abstract

Bovine leukemia virus (BLV), a retrovirus, causes Enzootic Bovine Leukosis (EBL) in cattle following a latent infection period. The BLV infection results in polyclonal expansion of infected B-lymphocytes and ∼5% of infected cattle develop monoclonal leukosis. Since the clonal expansion of virus-infected cell is a key in the pathogenesis of EBL, assessing the clonality of malignant cells is crucial for both understanding viral pathogenesis, which might be useful for EBL diagnosis.

For the investigation of clonality of BLV-infected cells in non-EBL and EBL cattle, two methods were used to evaluate the status of EBL; BLV-DNA-capture-seq method with high sensitivity and specificity and simple and cost-effective Rapid Amplification of Integration Site for BLV (BLV-RAIS) method. We found that the RAIS method efficiently detect expanded clone in EBL tissue sample as BLV-DNA-capture-seq method. Taking advantage of high frequency of BLV-infected cells in blood, we simplified RAIS method and showed that similar to BLV-DNA-capture-seq, this method could reliably provide quantitative value about clonal abundance of BLV-infected cells.

Next, we aimed to establish a diagnostic blood test for EBL by using the clonality information. First, we compared clonality of BLV-infected cells in blood with that in tumor tissue in EBL cattle. There was a remarkably similar clonality between blood and tissue in each animal. Furthermore, BLV integration site information clearly showed that the same clone was the most expanded in both blood and tumor tissue, indicating that tumor cells were circulating in blood in the disease cattle. We also analyzed tumor tissue at two independent anatomical regions and found the same clones was most expanded in both regions, supporting the idea that tumor cells are systemically circulating in the diseased cattle. Finally, we compared clonality value between non-EBL and EBL cattle by using BLV-RAIS method and found that there was clear difference between non-EBL and EBL. More importantly, we found that clonality value was low in asymptomatic phase but high in EBL phase in the longitudinal cohort study.

These findings have demonstrated that BLV integration site and clonality value are is a useful information to establish diagnostic blood test for EBL. That would contribute to reduction of economic damage caused by EBL and improvement of productivity in cattle industry.

## Introduction

Bovine leukemia virus (BLV) is a retrovirus that induces a life-long infection in a subset of B lymphocytes in cattle and causes Enzootic bovine leukosis (EBL) in some of the infected cattle after a long period of latency (1, 2). The integrated provirus provokes polyclonal expansion of the infected B lymphocytes and ∼5% of infected cattle develop monoclonal leukosis (3). Excluding Western Europe, worldwide approximately 50 million dairy cattle are BLV infected (4, 5). Due to the easy transmissibility of the virus, the infection is increasing hence increasing the risk of EBL onset in endemic countries. A high prevalence of BLV infection is reported in Japan, ∼40.9% and ∼28.7% in dairy and beef breeding cattle respectively (6). BLV infection leads to significant economic losses in the dairy farms in various countries (4, 7, 8).

BLV infection is confirmed by the presence of anti-BLV antibody in the blood. When an animal is in the leukosis stage, the disease is diagnosed by the presence of the tumors and/or general lymph node enlargement. The diagnosis of EBL generally depends on physical examination by a veterinarian. To reduce the variation among examiners, new objective and quantitative diagnostic tests are required. Since BLV is a retrovirus, the viral genome is integrated into the host genomic DNA. Considering that the huge size of the host genome, the viral integration sites are distributed to various location in the host genome and therefore be unique for each individual BLV-infected clone. One can use the viral integration site to distinguish individual infected cell. In asymptomatic phase of infection, there is a wide variation of infected clones observed in each individual cattle. During disease progression, certain clones are preferentially selected and expand clonally due to the host genome mutations which improves cell survival and/or cell proliferation. Finally, a certain clone becomes remarkably expanded and leads to EBL onset. A similar phenomenon is observed in the case of human T-cell leukemia virus type 1 (HTLV-1) infection where in the later stages, a single, monoclonal clone becomes the dominant clone and causes adult T-cell leukemia/lymphoma (ATL), a cancer of HTLV-1-infected T cell.

The application of next-generation sequencing (NGS) for clonality analysis is currently well established and provides an objective method to quantify the clonality of retrovirus-infected cells (9). Previous reports have showed that NGS is useful for the clonality analysis of BLV-infected cells (10–12); however, the NGS method is not feasible for practical use, especially for non-human retroviral infection like BLV. Several other molecular techniques which are mostly PCR-based have been reported to evaluate B-cell clonality or proviral load (13). However, these conventional methods are also not feasible for practical use to use as diagnose EBL.

Recently, a novel approach termed Rapid amplification of integration site (RAIS), has been developed to amplify HTLV integration sites, and followed by Sanger sequencing to assess the clonality of virus-infected cells (14). This method provides equivalent information as NGS-based methods at cheaper costs and shorter time frame. That motivated us to apply this method to the diagnosis of EBL. In this study, we modified the RAIS method to amplify BLV integration sites to develop a blood test for the diagnosis of EBL. We evaluated this method by analyzing non-EBL and EBL cattle including longitudinal samples collected from 2 to 3 years prior to the diagnosis of tumor onset.

## Result

### Establishment of “RAIS-Sanger sequencing approach” to assess the BLV clonality

The concept of rapid amplification of integration site (RAIS) method for BLV, termed BLV-RAIS, is shown in Fig. S1. There are various infected clones in asymptomatic (AS) cattle (12) and some cattle develop EBL after a long latency period. Genomic DNA was extracted from blood or tissue samples and analyzed by BLV-RAIS. As there are various clones in blood from AS cattle, the sequence next to the 3’LTR would be heterogenous. In contrast, the sequence would be homogenous in tumor from EBL cattle due to monoclonal expansion of the same clones. (Fig. 1A).

**Figure 1:**
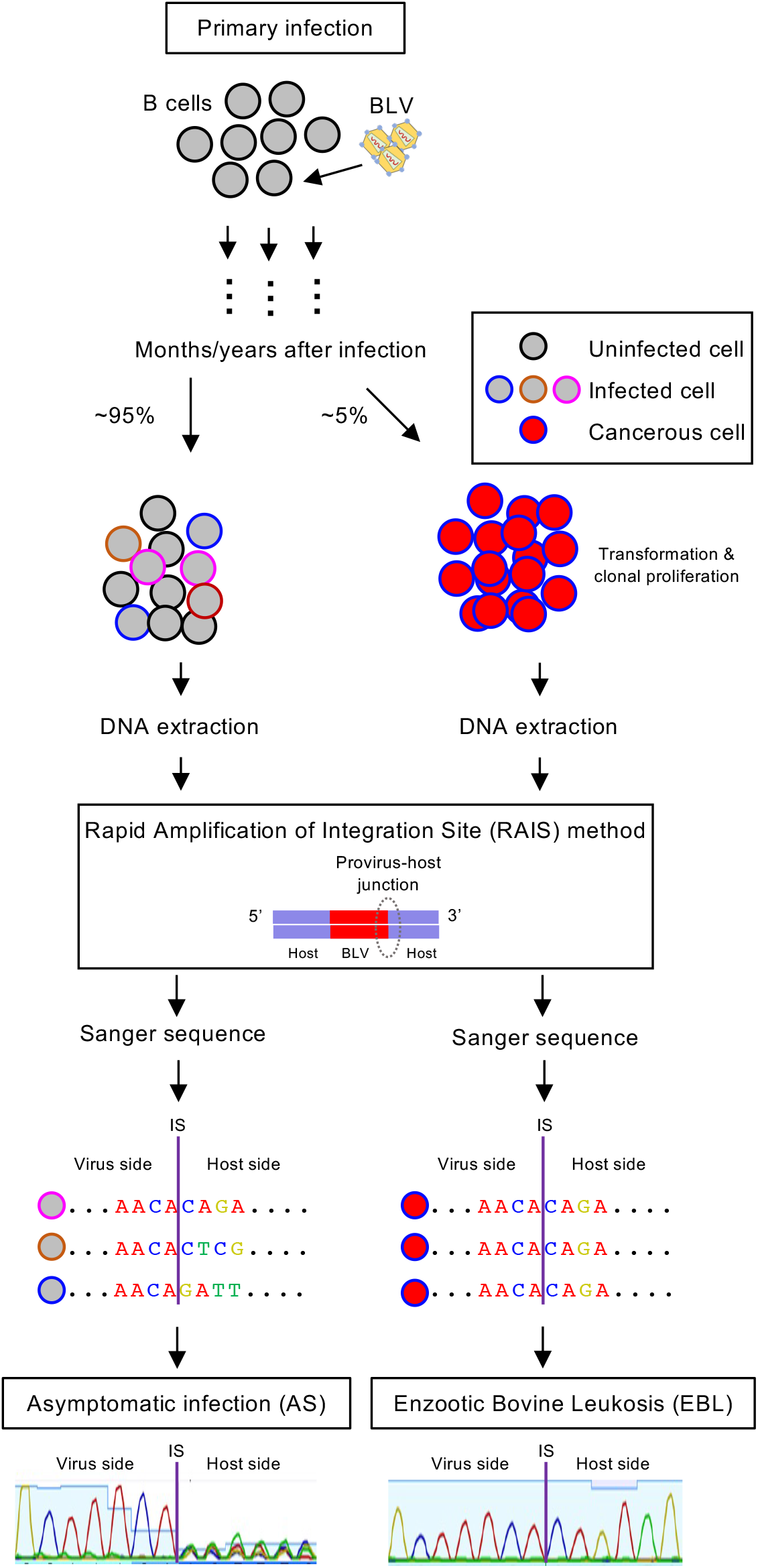
Schematic figure showing the identification of provirus clonality in BLV-infected B cells. Bovine leukemia virus (BLV) establishes latent infection by integrating the virus genome into host chromosomes of B cells and causes enzootic bovine leukosis (EBL) with monoclonal expansion of the infected cells. The different clinical stages of BLV infection are characterized by clonal distribution patterns of the provirus in the infected B cells. Rapid Amplification of Integration Site (RAIS) method has been utilized to amplify the 3’ end of the provirus-host junction. Due to variation in clone distribution of the infected B cells, Sanger sequencing chromatogram of the host side in the virus-host junction indicates a noisy and high background pattern in asymptomatic infection (AS) having polyclonal infected B cells. But in the case of EBL, the host and virus sides are equally clear and readable because of the monoclonal proliferation of the B cells.

The BLV-RAIS method was designed by modifying the original RAIS method for HTLV-1 (Fig. 1S)(14). First, the genomic regions containing the virus and the flanking host genome were amplified by ssDNA synthesis with biotinylated primer located just upstream of the 3’-LTR. Second, a polyA sequence was added to the 3’ end of ssDNAs containing virus-host junction followed by dsDNA synthesis with oligo dT primers. Next, we perform a nested PCR to further amplify the virus-host junction after Streptavidin purification. Finally, the PCR products were analyzed by Sanger sequencing. In order to establish a method which is applicable to a wide range of BLV genotypes, the primers for BLV-LTR should be designed at conserved regions among various cattle samples. We analyzed the LTR sequence by using DNA-capture-seq data obtained previously (15) and LTR sequences available in public database. We then designed primers in the BLV-LTR that match various genotypes of BLV (Fig. S2).

To provide proof of concept for the BLV-RAIS method, we first analyzed eight BLV-infected cattle in which BLV clonality was determined by viral DNA-capture-seq analysis (15) (Fig. 2A). Summary of the characteristics of eight cattle are shown in the Table S1. We analyzed the BLV-RAIS products by agarose gel electrophoresis and found there was smear in each sample, suggesting we were able to amplify the BLV integration sites using our designed primers (Fig. 2B). To confirm that, we performed Sanger sequencing analysis using the primer located in the BLV-LTR to obtain DNA sequence containing the virus-host junction. The obtained sequences were aligned against the reference BLV-LTR sequence (GenBank: EF600696.1) for identification of the virus-host junction. The DNA sequence of virus region was clear; however, the flanking host genome sequence next to the end of BLV-LTR were heterogenous (Fig. 2C), suggesting that there was no significantly expanded clone. The DNA sequence of the flanking host genome were heterogenous but contained some dominant fluorescent peaks in PL2 and PL3 samples (Fig. 2C), suggesting the presence of some expanded clones. In contrast, the sequencing results from EBL cattle were homogenous both in viral and the flanking host region, suggesting the presence of a single dominant clone (Fig. 2D). These observations were consistent with the clonal distribution pattern obtained by the NGS analysis (Fig. 2A).

**Figure 2:**
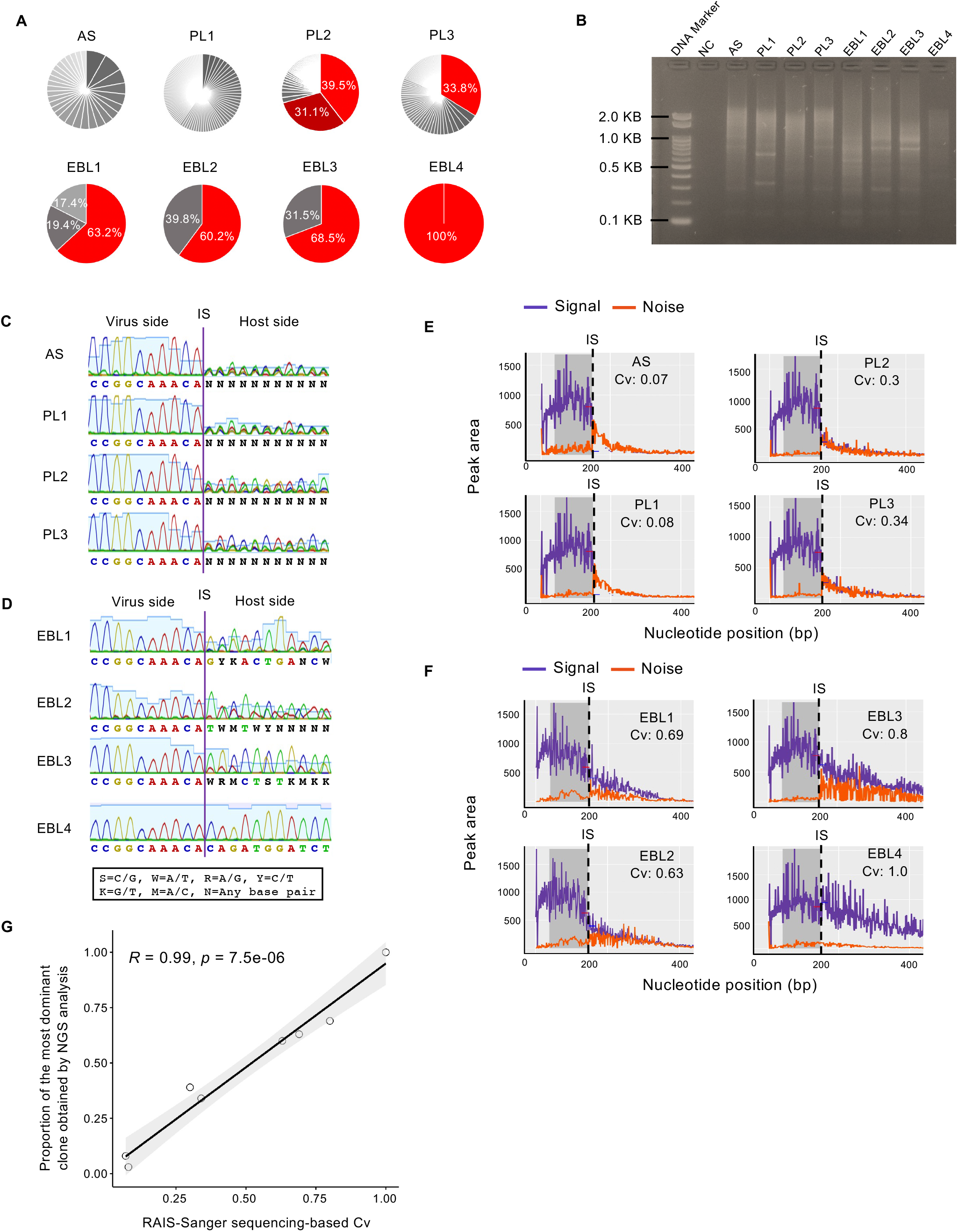
BLV-RAIS method with sanger sequencing reveals BLV clonality in cattle having different clinical features. (A) Clonal abundance pattern of eight BLV-infected cattle. PBMCs of one asymptomatically infected (AS) and three persistent lymphocytosis (PL) cattle and tumors of four EBL cattle were analyzed by conventional RAIS method and next generation sequencing (NGS) to compare the clonality of BLV-infected cells. Each pie chart represents the clonal distribution for an individual sample and each slice represents an individual clone. (B) Gel image of BLV-RAIS-amplified products of 3’ end of the virus-host junction from 500 ng of sample DNA. DNA from BLV-uninfected cattle was used as negative control. NC: negative control. (C and D) Sanger sequencing chromatograms of RAIS-amplified products from virus-host junction of AS and PL (C) and EBL (D) cattle. IS: Integration Site. (E and F) CLOVA analysis of Sanger sequencing data for comparison of signal-noise chromatograms which also automatically calculated the clonality value (Cv). (G) Correlation analysis between the Sanger sequencing-based Cv and the proportion of the most dominant clone obtained from NGS analysis (Pearson correlation test) for samples of the eight BLV-infected cattle.

To quantify the clonality of BLV infected cells, the CLOVA software was used (16). Based on the Sanger sequence spectra, CLOVA generates a signal-noise chromatogram and automatically performs an analysis to express the clonality as the Clonality value (Cv) (Fig. 2E and 2F). The Cv ranges between 0 and 1 where 0 indicates that infected clones in the sample are completely polyclonal whereas 1 indicates monoclonal. We compared the Cv and the proportion of the most dominant clone obtained from NGS analysis and observed a strong correlation between the two values (Fig. 2G). These findings demonstrated that the BLV-RAIS can quantify the clonality of BLV-infected cells at similar accuracy as the NGS-based assay.

### Modification of BLV-RAIS to simplify the protocol and its representativeness

To increase the feasibility of the method so that it can be used as a routine diagnostic method for EBL, we performed some modifications to simplify the BLV-RAIS protocol. The original RAIS method was developed for clonality analysis of HTLV-infected cells, in which the proviral load (PVL) is generally low in asymptomatic carriers. The PVL of BLV is higher than that of HTLV-1 (Fig. 3A), which suggests that we would be able to skip the Streptavidin bead purification step (Fig. 3B), thus simplifying the protocol and making it much more time and cost-effective. We then analyzed six BLV infected cattle with various clonality using both the conventional and the modified BLV-RAIS method. There was a strong correlation between the Cv obtained from conventional and modified BLV-RAIS method (Fig. 3C) suggesting that skip of the Streptavidin bead purification step does not change the robustness of the BLV-RAIS method to quantify clonality of BLV-infected cells. Next, we analyzed how reproducible the BLV-RAIS method is within technical replicates and found little to no difference (Fig. 3D). We further tested the variation of the Cv when we analyzed the same samples at different institutes and examiners. We found a strong linear correlation between the Cv of the independent analysis (Fig. 3E). These data showed that BLV-RAIS method could provide reproducible value even when the analysis was performed at different institutes by different examiners.

**Fig. 3.**
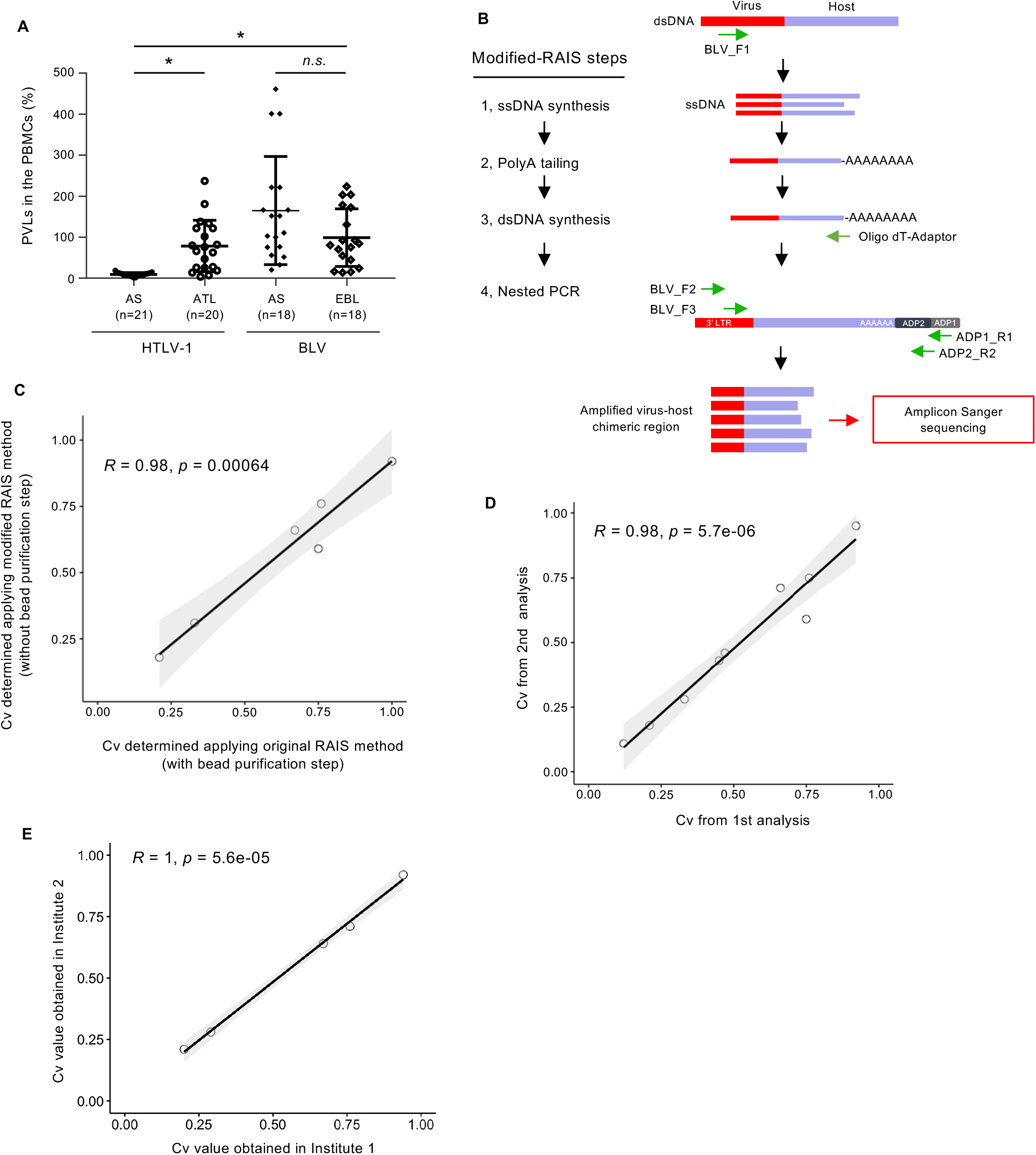
New RAIS method with simplified protocol and its feasibility. (A) Comparison of proviral loads (PVLs) of HTLV-1-infected patients and BLV-infected cattle. PVLs indicate the number of virus-infected cells per 100 PBMCs. The PVL of HTLV-1 was referred from our previous study (21). The bars indicate median values with a 95% confidence interval. Statistical significance was obtained by the Mann-Whitney U test and Kruskal-Wallis test. *: p<0.0001; *n.s.*, not significant. (B) Schematic figure of modified RAIS method for detection of BLV clonality. The conventional RAIS method has an avidin-bead purification step to enrich virus-host chimeric double-stranded DNAs linked with dT-adaptor (Fig. S1). Considering the characteristic high PVL in BLV infection, we developed a modified-RAIS method without the purification process. (C-E) Correlation analysis (Pearson correlation test) of the Cv to compare modified and conventional BLV-RAIS methods (C), representative analysis (D), and same samples analyzed in two independent institutes (E).

### Clonality of BLV-infected cells in the blood is well correlated with that in tumor tissue in EBL cattle

In Fig. 2 and Fig. 3, we analyzed tumor tissues from EBL cattle and blood samples from non-EBL cattle. To apply the BLV-RAIS method for EBL diagnosis even before slaughtering the cattle, it is vital to ascertain if we can use peripheral blood samples to predict the presence of tumor tissue in the cattle. To achieve this, we compared the clonality of BLV-infected cells between blood and tumor tissue from the same EBL cattle. First, we quantify the clonality of BLV-infected cell in the DNA samples from 11 EBL cattle (Table S2) by using viral DNA-capture-seq method. The clonal distribution in the tumor tissue and the peripheral blood were quite similar and the identical dominant clone was found in both the tissue and peripheral blood at almost identical abundance obtained by NGS-based method (Fig. 4A). These data have indicated that tumor cells are systemically circulating in EBL cattle by blood flow. Consistent with the idea, the same dominant clone was detected in two independent tumor tissues at different anatomical locations (Table S3, Fig. 4B). Next, we evaluated efficiency of BLV-RAIS by analyzing blood samples. As shown in Fig. 2G, we found that Cv were consistent with the clonal abundance of the most expanded clone in peripheral blood (Fig. 4C). There was also a strong linear correlation between the Cv of blood and tissue (Fig. 4D). These findings demonstrated that the BLV-RAIS method can be a useful blood test for diagnosis of EBL.

**Fig. 4.**
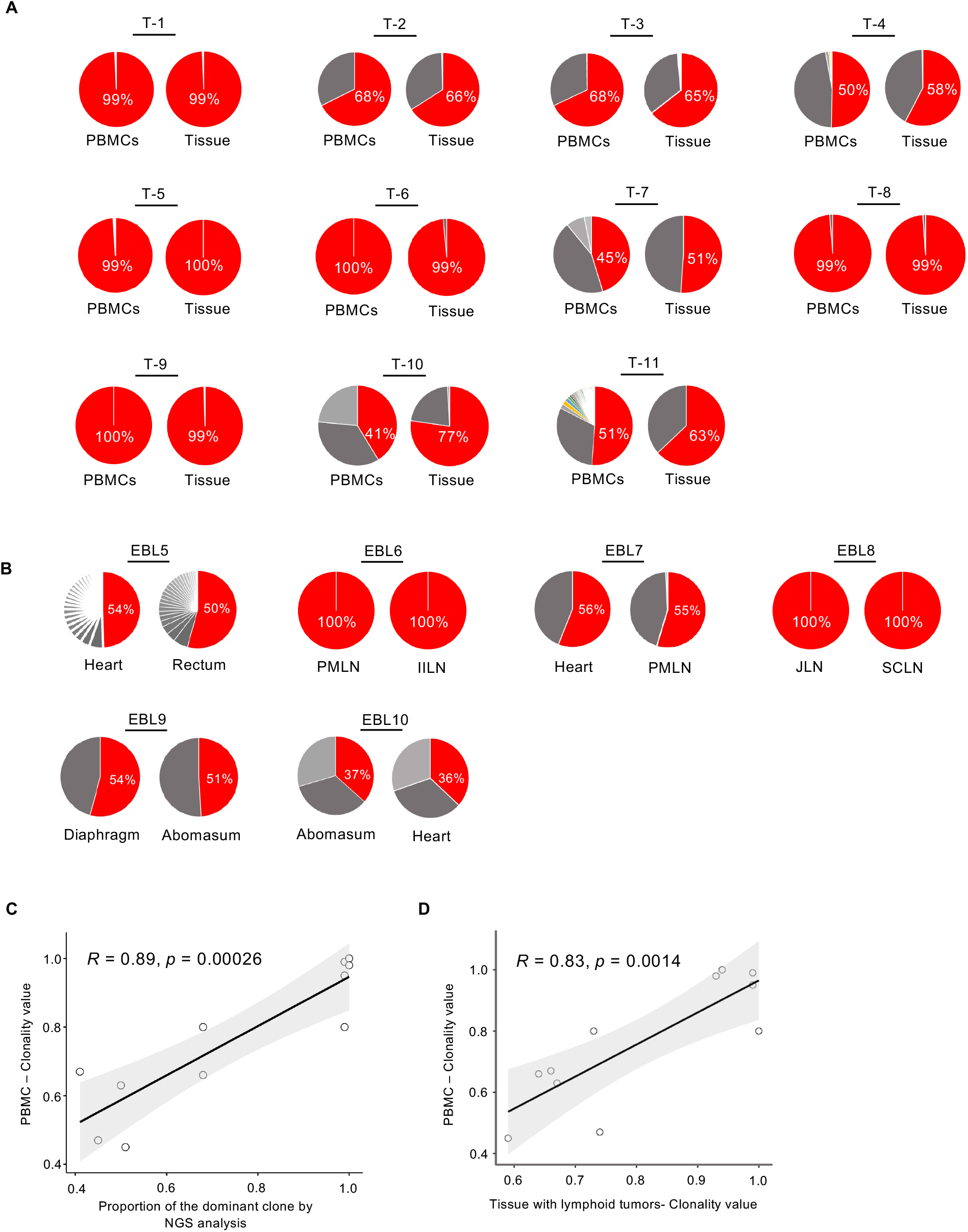
Comparison of BLV clonality in PBMCs and tumor tissues in EBL cattle by using modified RAIS method. (A and B) Clonal abundance distribution of BLV in PBMCs/tissues with lymphoid tumors (A) and paired two different tissues (B) from EBL cattle. BLV-host junction in infected cells was amplified by modified RAIS methods. PMLN: Posterior mediastinal lymph node, IILN: Internal iliac lymph node, JLN: Jejunal lymph node, SCLN: Superficial cervical lymph node. (C) The proportion of dominant BLV clones was plotted against the BLV-RAIS Sanger sequencing-based clonality value. (D) Correlation analysis (Pearson correlation test) between the BLV-RAIS Sanger sequencing-based Cv obtained from the PBMCs and tissues of EBL cattle. The proportion and Cv of PBMCs in Fig. 4C and D were derived from eleven cattle shown in Fig. 4A.

### Analysis of longitudinal samples to test the possibility of identifying pre-EBL cattle

We further investigated whether the BLV-RAIS can distinguish non-EBL and EBL status in the cattle in a longitudinal cohort study. We analyzed five BLV infected cattle that develop EBL and had longitudinal samples at two different time-point and at the time of EBL diagnosis (Table 1). The time gap between the second and initial sampling time was over two years in all five cases and between these two-time points, the PVL increased in three cattle while decreased in the other two cattle (Table 1). We performed viral BLV-DNA-capture-seq using all the longitudinal samples to characterize when the dominant clone in tumor was detected and how the clone expanded in longitudinal samples (Fig. 5A). There were two expanded clones in C1 sample while the other 4 cases contained a single expanded clone. Among the five EBL cases, the dominant BLV-infected clones in the tumor were undetectable in the longitudinal blood samples of two cattle, C2 and C5. In cattle C2, the EBL clone was detected at 71 days before the diagnosis of tumor onset with approximately 44% occupancy. The EBL clone was detectable even at 869 days before tumor onset. In cattle C5, the dominant integration site in tumor was detected in the blood at 245 days before tumor diagnosis but the frequency was low (2.7%). The Cv of each sample were well correlated with the degree of the most expanded clone in EBV-DNA-capture-seq. We also performed BLV-RAIS and found Cv was high in all EBL cattle but low in non-EBL cattle, including blood sample obtained at 71 days before the diagnosis of tumor in cattle C2.

**Figure 5.**
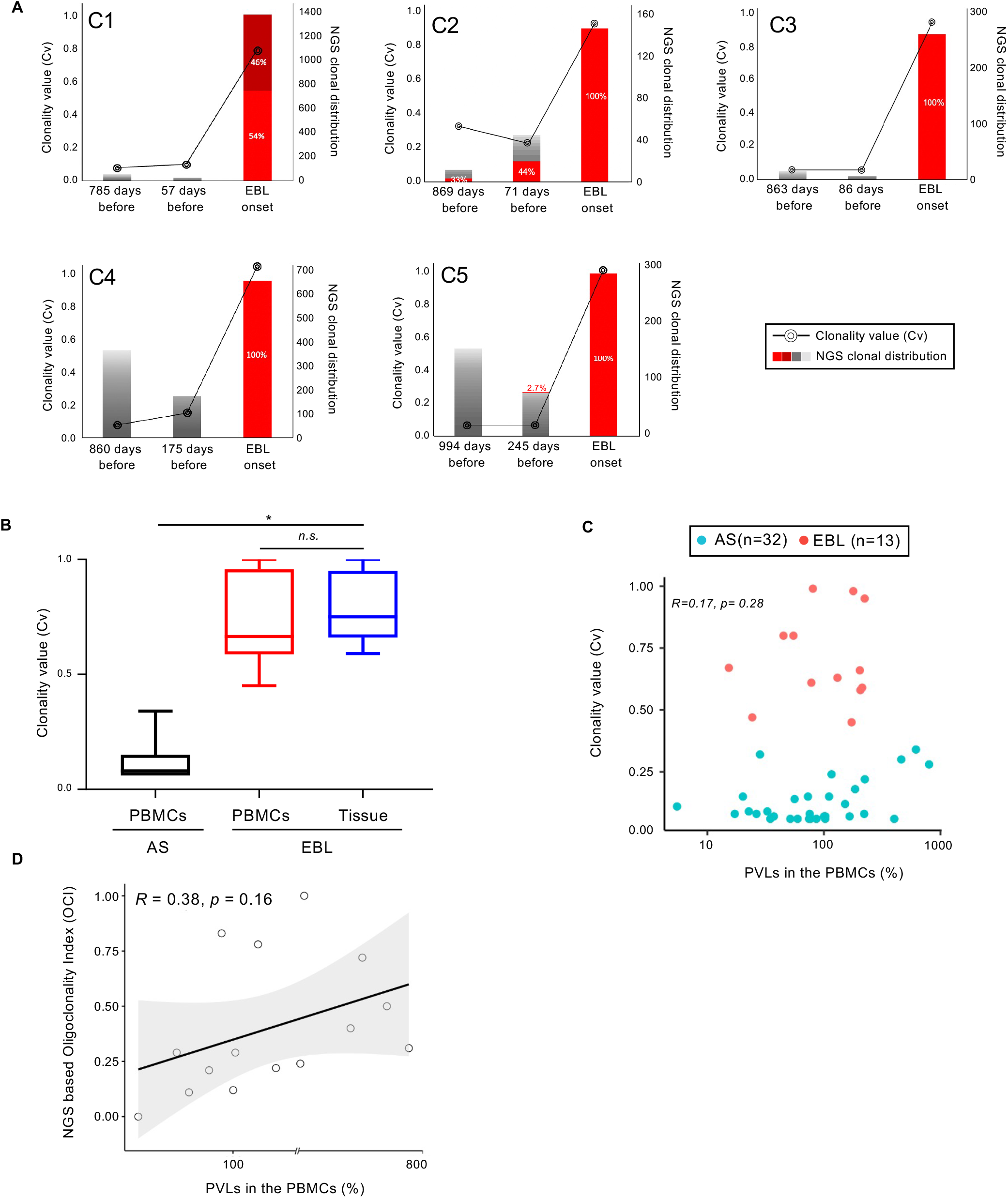
Tracking of tumor-dominant BLV integration sites before diagnosis of EBL onset. (A) BLV integration sites in PBMCs and tumor biopsies (EBL onset) from five cattle were quantitatively mapped by high throughput sequencing. The line graph depicts the Cv obtained from modified BLV-RAIS method at selected time points (data plotted on the left Y-axis). The number of BLV integration sites is shown as bar graph (data plotted on the right Y-axis). Gray and red sub-bars represent unique and tumor-dominant BLV integration sites as shown NGS clonal distribution, respectively. (B) Comparison between the Cv range of asymptomatic (AS) and EBL cattle. Statistical significance was obtained by the Mann-Whitney U test and One-way ANOVA, 95% confidence interval; *:p<0.0001, *n.s.*, not significant. (C) Cv obtained from PBMCs of BLV-infected cattle were plotted against the PVLs in PBMCs. (D) Correlation analysis (Pearson correlation test) between the PVLs in the PBMCs of BLV-infected cattle and NGS based Oligoclonality Index (OCI).

**Table 1:** Summary of EBL cohort used in this study

| Cattle ID. | Time point 1 |  | Time point 2 |  | EBL |
| --- | --- | --- | --- | --- | --- |
|  | Days before EBL diagnosis | PVL in PBMC (%) | Days before EBL diagnosis | PVL in PBMC(%) | Name of tissue tumor site used in this study |
| C1 | 785 | 100.27 | 57 | 32.68 | Diaphragm |
| C2 | 869 | 28.26 | 71 | 224.35 | Heart |
| C3 | 863 | 74.6 | 86 | 102.6 | Abomasum |
| C4 | 860 | 102.9 | 175 | 56 | Lymph node |
| C5 | 994 | 66.6 | 245 | 86.1 | Heart |

In this study, we’ve analyzed 78 blood and tumor samples from 42 BLV-infected cattle. The Cv obtained from PBMC and tumor samples of the EBL cattle ranged from 0.45 to 1.0 which was significantly higher than the Cv range of asymptomatic cattle (p-value <0.01) (Fig. 5B). We plotted the PVL and the Cv obtained from PBMC and found there was no significant correlation (Fig. 5C). Similar to the Cv, we didn’t find a significant correlation between the NGS-based oligo clonality index (OCI) and the PVL in the PBMC (Fig. 5D), indicating that PVL is partially but not perfectly reflect the status of EBL progression.

## Discussion

The incidence of EBL is increasing in some endemic country such as Japan. This induce a serious problem for the farm to maintain cattle. Current diagnosis of EBL heavily depends on physical examination by a veterinarian. Once there are abnormal findings of obvious enlargement of superficial lymph nodes, or organs palpable by rectal examination, the EBL-suspected cattle is subjected to further examination of blood smear test to detect atypical lymphocyte and/or histological test. There is a risk of underdiagnosis especially when the EBL-related tumor is still not at the advanced stage. Therefore, we need to develop some objective, quantitative and highly-sensitive diagnostic test for tumor of EBL. Several candidate markers detected using blood tests has been reported to be useful to diagnose EBL. Serum total lactate dehydrogenase (LDH) and thymidine kinase (TK) tends to be elevated in EBL cattle (17). Since EBL is a B-cell-lymphoma, immunophenotypic analysis of peripheral blood is also reported as a candidate blood test (18). These markers might be helpful to diagnose EBL; however, they still have limited specificity as LDH and TK levels may be elevated in other type of cancers. Immunophenotypic analysis can be affected by some inflammatory diseases. EBL is a cancer caused by BLV; therefore, the analysis of BLV would be the most specific and sensitive way to evaluate the status of EBL. PVL is a value obtained by quantification of viral and the host genomic DNA copy. PVL has been used to evaluate the proportion of infected cells in blood or tumor tissue. We and others have showed that PVL is not specifically elevated in EBL. Asymptomatic and non-EBL cattle showed high proviral load as EBL cattle (Fig. 3A)(17). Thus, it is not so useful to know the quantity of infected cells but the clonality of infected cells is critical to know the disease status of EBL (Fig. 5B). This seems logical because clonal expansion of BLV-infected cells is directly associated with oncogenesis of EBL. Thus, the diagnostic test based on oncogenesis of BLV-infected cells, namely extent of clonal expansion, will provide the highest specificity.

Clonality analysis of retrovirus-infected cells was drastically changed by application of NGS in HTLV-1 infection (9). The method is applied for other retroviruses, such as HIV-1 (19, 20) and BLV (10–12), resulting in better understanding of *in vivo* dynamics of retrovirus-infected cells with high resolution and accuracy. However, these methods are not suitable for practical use because of its cost and technical requirement to perform. The protocol to use rapid amplification of integration site followed by Sanger sequencing was reported by Saito et al to quantify clonality of HTLV-1-infected cells (14). That led us to apply the method for BLV infection.

EBL is evident in tumor tissue, but we need to confirm if tumor cells are also present in peripheral blood when we apply the RAIS method as a blood test for EBL. To answer the question, we analyzed 11 paired DNA samples, blood and tumor tissue, obtained from each the same EBL cattle. The result clearly demonstrated that the same expanded EBV-infected clone was detected in blood as well as in the tumor tissue with equivalent dominance in all 11 EBL cattle we analyzed (Fig. 4A). To our knowledge, this is the first report to provide concrete evidence that the same tumor cells in lymphoma tissues are circulating in peripheral blood in EBL cattle. This key evidence further suggested the possibility to apply RAIS method as a blood test for EBL. Consistent with a recent study which also demonstrated that biotin-streptavidin purification step can be skipped both in HTLV-1 and BLV samples (16), we simplified the RAIS protocol to increase the feasibility for practical use by removing the biotin-streptavidin purification step (Fig. 2B and 2C). In this study, we obtained the evidence that BLV-RAIS method would be a potent blood test to diagnose EBL. It has been proposed that early detection of ATL might be possible as the tumor dominant integration site started to evolve years before the onset of symptoms (5). We also showed that the same EBL clone was detectable before the onset of the disease in two of five EBL cattle in longitudinal study (Fig. 5A). However, there are no detectable EBL clone even just 2 to 6 months before the disease onset in three of five cattle. Further study with large number of cases is required to know whether BLV-RAIS is useful for the early diagnosis of EBL.

In summary, we provided the evidence that BLV-RAIS can be a quantitative and reproducible blood test useful for the diagnosis of EBL. Further investigation would be required to maximize the advantage of BLV-RAIS method to monitor the EBL, improve productivity of cattle industry and contribute to establishment of Sustainable Development Goals.

## MATERIALS AND METHODS

### Regulatory Approvals

This study was approved by the animal research committee at Tokyo University of Agriculture for sample collection at farms. We confirm that all experiments were performed in accordance with the committee’s guidelines and regulations. The farm leaders gave verbal consent for the blood sample collection, serological test of BLV antibodies, and PVL measurements. All data or information excludes personally identifiable information such as farmers’ names.

### Collection and storage of the peripheral blood and tissue biopsies

We utilized excess blood and the serum samples collected by Kanagawa Shonan Livestock Hygiene Service Center from 2015 to 2019 for national whole herd test surveillance targeting Johne’s disease. The blood samples were stored at −20℃ until analysis. The visually identifiable lymphoma tissues for the cohort study (Fig. 5) were collected from the carcasses after the postmortem inspections at Meat Inspection Station, Kanagawa Prefectural Government. The peripheral blood and visually identifiable lymphoma tissues for comparative study (Fig. 4) were collected from the carcasses after the postmortem inspections at Kumamoto Prefectural Meat Safety Inspection Office from December 2021 until March 2022. The blood samples were stored at 4℃ and lymphoma tissues were kept at −20℃ until analysis.

### Separation of peripheral blood mononuclear cells (PBMCs) and extraction of genomic DNA from PBMC and tissue biopsies

Peripheral blood was processed by density-gradient centrifugation using Ficoll-Paque (GE Healthcare) for the separation of PBMCs and cryopreserved at −80 °C in Cell Banker (Juji Field Inc.) until use. For the extraction of genomic DNA from whole blood, PBMCs, and tissue biopsies, the DNeasy Blood and Tissue kit (Qiagen) was used according to the manufacturer’s protocol.

### Quantification of proviral load (PVL) in PBMC and tissue biopsies

The PVLs in PBMC and tissue biopsies were estimated by quantifying the number of copies of the BLV pro gene and bovine beta-actin gene using the digital droplet PCR as described previously (21). The PVL was expressed in copy per 100 cells as follows, PVL (%) = [(copy number of BLV pro)/ (copy number of bovine beta-actin)/2] × 100. Primer sequences are listed in Table S4.

### DNA capture seq. for mapping and quantification of BLV integration sites (IS)

BLV DNA-capture-seq and mapping of viral integration sites were performed as previously described (15, 21, 22) with minor modifications. Briefly, 2 μg genomic DNA was sonicated using a Picoruptor (Diagenode s.a., Liège, Belgium) to produce 300–500-bp fragments. DNA libraries were prepared using a NEBNext Ultra II DNA Library Prep Kit and NEBNext multiplex Oligos for Illumina (New England Biolabs). BLV-specific probes were used for the enrichment of DNA libraries with BLV sequence and the enriched libraries were PCR amplified using the primer set P5-P7. The enriched DNA libraries were quantified by the TapeStation instrument (Agilent Technologies) before sequencing via Illumina MiSeq or NextSeq.

The three FASTQ files (Read 1, Read 2, and Index Read) obtained from the Illumina sequencing went through a data-cleaning step and the cleaned sequences were aligned to *Bos taurus* reference genome sequence (ARS-UCD1.2/bosTau9) with BLV (GenBank: EF600696) as a separate chromosome or integrated provirus using the BWA-MEM algorithm (23). The aligned reads were visualized by Integrative Genomics Viewer (IGV) (24) after some additional processing and clean-up using Samtools (25) and Picard (http://broadinstitute.github.io/picard/). For IS analysis, we aligned the cleaned FASTQ files to the reference genome containing bovine chromosomes (Chr 1–29 and X) and BLV as 2 separate chromosomes: the whole viral sequence excluding the LTRs (BLV_noLTR) and viral LTR sequence as a separate chromosome (BLV_LTR). The chimeric reads containing both the BLV and host genome were extracted to identify the locations of BLV IS in the host genome using the method described previously(15, 21). The number of final virus-host reads in a certain genomic region reflects the initial cell number for each infected clone that enabled us to estimate the clonal abundance for each infected clone (9).

### Amplification of BLV-host chimeric region by RAIS method

Amplification of the BLV-host chimeric region was performed via the conventional and modified RAIS method using the BLV specific forward primers: BLV_F1-TCTCCCGCCTTTTTTGAGG for the single-strand DNA synthesis, BLV_F2-CAGACCTTCTGGTCGGCTATC, and BLV_F3-CGAGCTCTATCTCCGGTCCT for the final nested PCR amplification. The BLV_F1 primer binding site is at a position slightly upstream of the 3’ LTR and the binding sites for BLV_F2 and BLV_F3 are at the 3’LTR (Fig. S2). The detailed protocol is provided as supplemental method. The persistently BLV-infected fetal lamb kidney cell line (FLK-BLV) DNA was used to optimize the RAIS method for the BLV study. FLK-BLV cell line was established by Van Der Maaten and Mille (26) and was kindly provided by Dr. Yoko Aida, Institute of Physical and Chemical Research (RIKEN), Japan.

The RAIS amplified BLV-host chimeric regions were then purified using the QIAquick^R^ PCR purification kit (QIAGEN) and Sanger sequenced by a commercial DNA sequencing service (FASMAC Co., Ltd.).

### Quantification of BLV clonality using the CLOVA software

The Sanger sequence data of the virus-host chimeric region were analyzed using CLOVA (https://fasmac.co.jp/en/rais) providing 20 nucleotide sequences of the 3’ end of BLV LTR as the transgene sequence (16). CLOVA is a dedicated R program-based software that utilizes the “Sanger ab1 file” to analyze transgene sequences. Compared with the transgene, CLOVA generates a signal-noise chromatogram for the whole sequence and automatically calculates the clonality value (Cv) taking into consideration the value of signal peak area (intensity) of the host side and the virus side. In this study, signal peak areas up to 20 nucleotide positions upstream and downstream from the virus-host integration site were considered to calculate the Cv using CLOVA.

The Cv ranges between 0 and 1; 0 indicates host genome sequence is a contribution of multiple clones with an equal proportion of the load, and 1 indicates the host genome sequence is attributed by a single clone that dominates completely.

### Statistical analysis

All data analyses were performed using GraphPad Prism 7 software (GraphPad Software, San Diego, CA) and R (v4.0.3). Correlation analysis and the scatter plots were generated in R using the Pearson correlation test. The oligoclonality index (OCI) based on the Gini coefficient was computed using the Ineq R package (http://CRAN.R-project.org/ package5ineq) in R. The OCI is used to measure the non-uniformity of the clonal distribution which ranges between 0 and 1, with 0 indicating perfect polyclonal and 1 indicating perfect monoclonal. Because of the variable clone population, a type 1 correction was considered as described previously (9).

## Acknowledgements

We would like to express our gratitude to the staff of the Kumamoto Prefectural Meat Inspection Office and Meat Inspection Station, Kanagawa Prefectural Government for providing tumor samples of EBL cattle. We thank the staff of Kanagawa Shonan Livestock Hygiene Service Center for blood sample collection of BLV infected cattle. This work was supported in part by JSPS KAKENHI Grants-in-Aid for Scientific Research C 21K05943 (to T.K.) and Livestock Promotional Funds of Japan Racing Association (JRA).

## Authors’ contributions

T.K. and Y.S. conceived and coordinated the project; H.B., S.M., A.R., S.A.R. and N.O. performed experiments. H.B. and N.O. performed the bioinformatics analyses. B.H., T.K., S.M., M.M., K.N., BJY.T., K.S. and Y.S. analyzed the data. B.H., T.K., K.I. and Y.S. wrote the manuscript. T.I. and K.I. contributed to the materials and results interpretation. M. S. provides analytic tools. All authors read and approved the final manuscript.

## Additional Information

Competing financial interests: The authors declare no competing financial interests.

---

**Fig S1:**
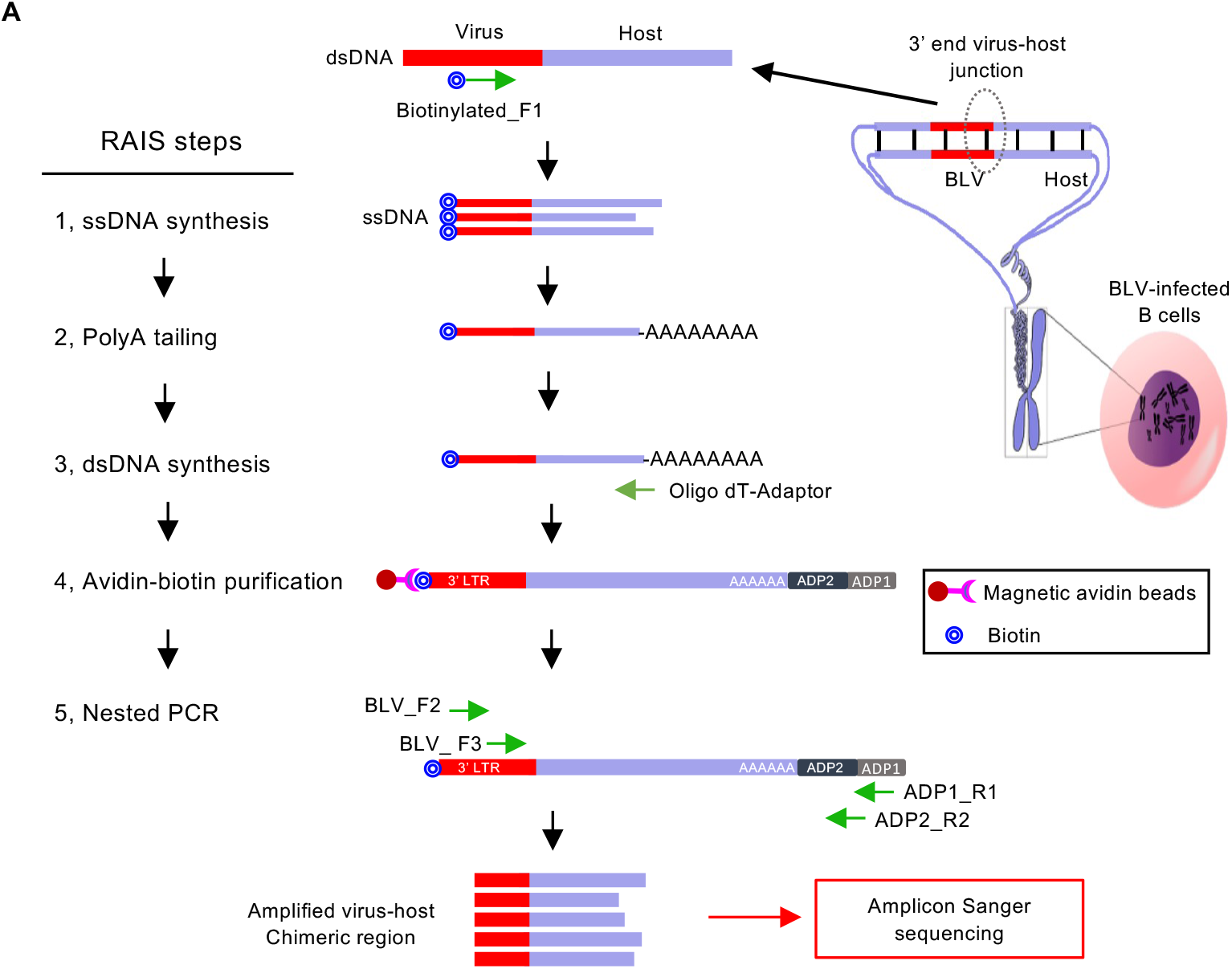
Schematic illustration of the different steps of BLV-RAIS method before modification.

**Figure S2.**
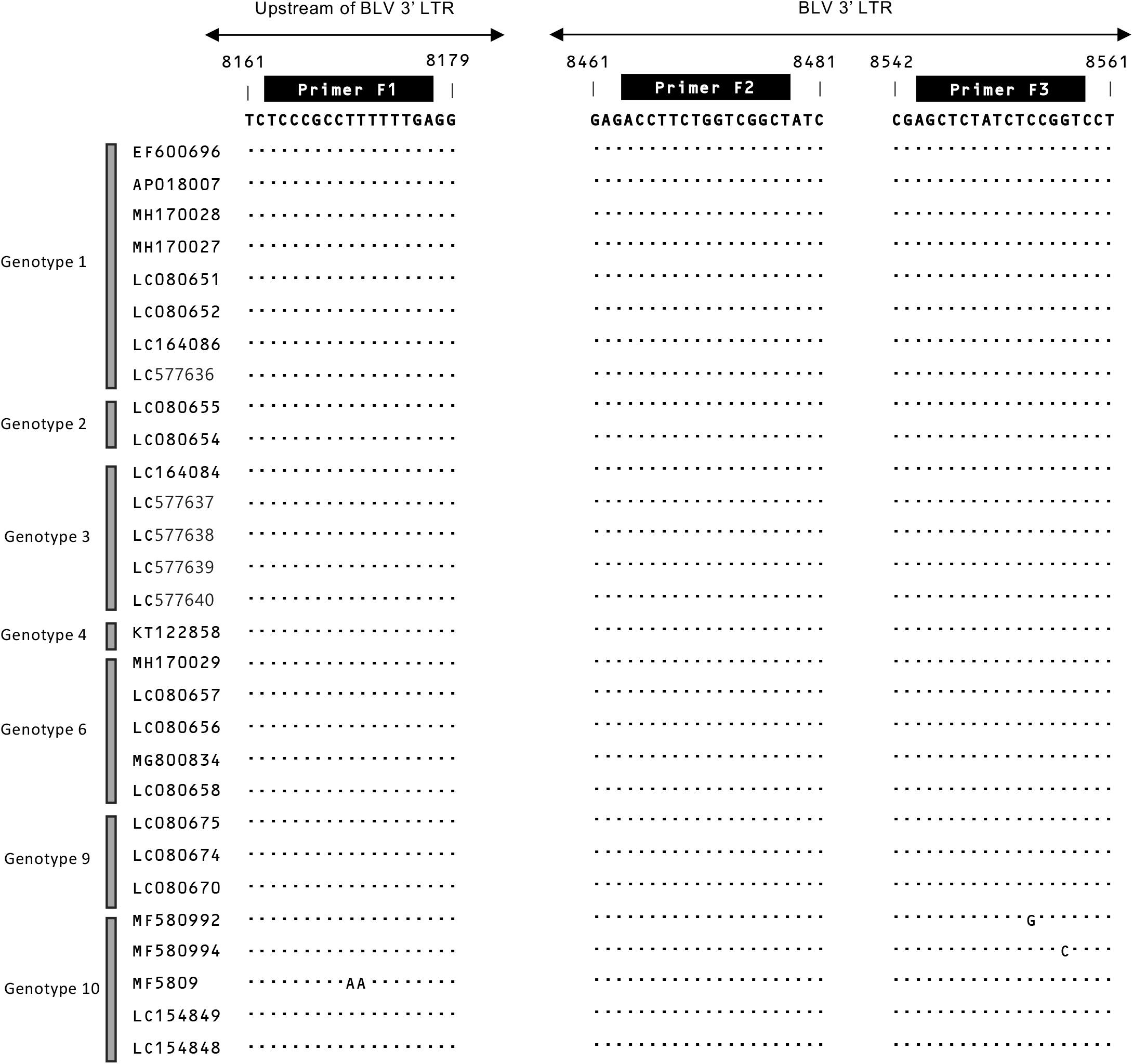
Location of RAIS primers in conserved regions of Bovine leukemia virus (BLV). Identity with BLV strain EF600696 is indicated by a dot. The FI, F2, and F3 primer regions are indicated on top of the alignment. There are no available LTR sequences of Genotypes 5,7 and 8.

**Table S1:** Summary of characteristics of cattle analyzed by conventional BLV-RAIS method.

| Disease group | Cattle ID. | Clinical status | PBMCs($\times 10^3/\mu\text{l}$) | Tissues with lymphoid tumors |
| --- | --- | --- | --- | --- |
| Asymptomatic | AS | Asymptomatic | 6.5 | - |
|  | PL1 | Persistent lymphocytosis | 12.8 | - |
|  | PL2 | Persistent lymphocytosis | 10.6 | - |
|  | PL3 | Persistent lymphocytosis | 13.4 | - |
| Enzootic Bovine Leukosis (EBL) | EBL1 | Enzootic bovine leukosis | NA | <u>heart</u>, kidneys, lung, rumen, uterus, intestine, lymph nodes |
|  | EBL2 | Enzootic bovine leukosis | NA | <u>heart</u>, lung, rumen, uterus, lymph nodes |
|  | EBL3 | Enzootic bovine leukosis | NA | <u>heart</u>, kidneys, lung, rumen, uterus, intestine, lymph nodes |
|  | EBL4 | Enzootic bovine leukosis | NA | <u>heart</u>, kidneys, rumen, uterus, intestine, lymph nodes |

underlined tumor samples were analyzed in this study; NA: not available; -: not applicable

**Table S2:** EBL cattle for which both the PBMC and tumor samples were analyzed.

| Cattle ID. | Clinical status | Liquid biopsy | Solid biopsy | Cattle ID. | Clinical status | Liquid biopsy | Solid biopsy |
| --- | --- | --- | --- | --- | --- | --- | --- |
| T-1 | EBL | PBMC | Uterus | T-7 | EBL | PBMC | Heart |
| T-2 | EBL | PBMC | 3<sup>rd</sup> stomach | T-8 | EBL | PBMC | 4<sup>th</sup> stomach |
| T-3 | EBL | PBMC | Mesenchymal lymph node | T-9 | EBL | PBMC | Thymus region |
| T-4 | EBL | PBMC | Iliac lymph nodes | T-10 | EBL | PBMC | Kidney lymph node |
| T-5 | EBL | PBMC | Heart | T-11 | EBL | PBMC | Kidney |
| T-6 | EBL | PBMC | Mediastinal lymph node |  |  |  |  |

**Table S3:** EBL cattle for which two different tissue tumor sites were analyzed.

| Cattle ID. | Clinical status | Tissues with lymphoid tumors |  |
| --- | --- | --- | --- |
|  |  | Tissue site-1 | Tissue site-2 |
| EBL5 | EBL | Rectum | Heart |
| EBL6 | EBL | Internal iliac lymph node (IILN) | Posterior mediastinal lymph node (PMLN) |
| EBL7 | EBL | Heart | Posterior mediastinal lymph node (PMLN) |
| EBL8 | EBL | Jejunal lymph node (JLN) | Superficial cervical lymph node (SCLN) |
| EBL9 | EBL | Diaphragm | Abomasum |
| EBL10 | EBL | Heart | Abomasum |

**Table S4.**
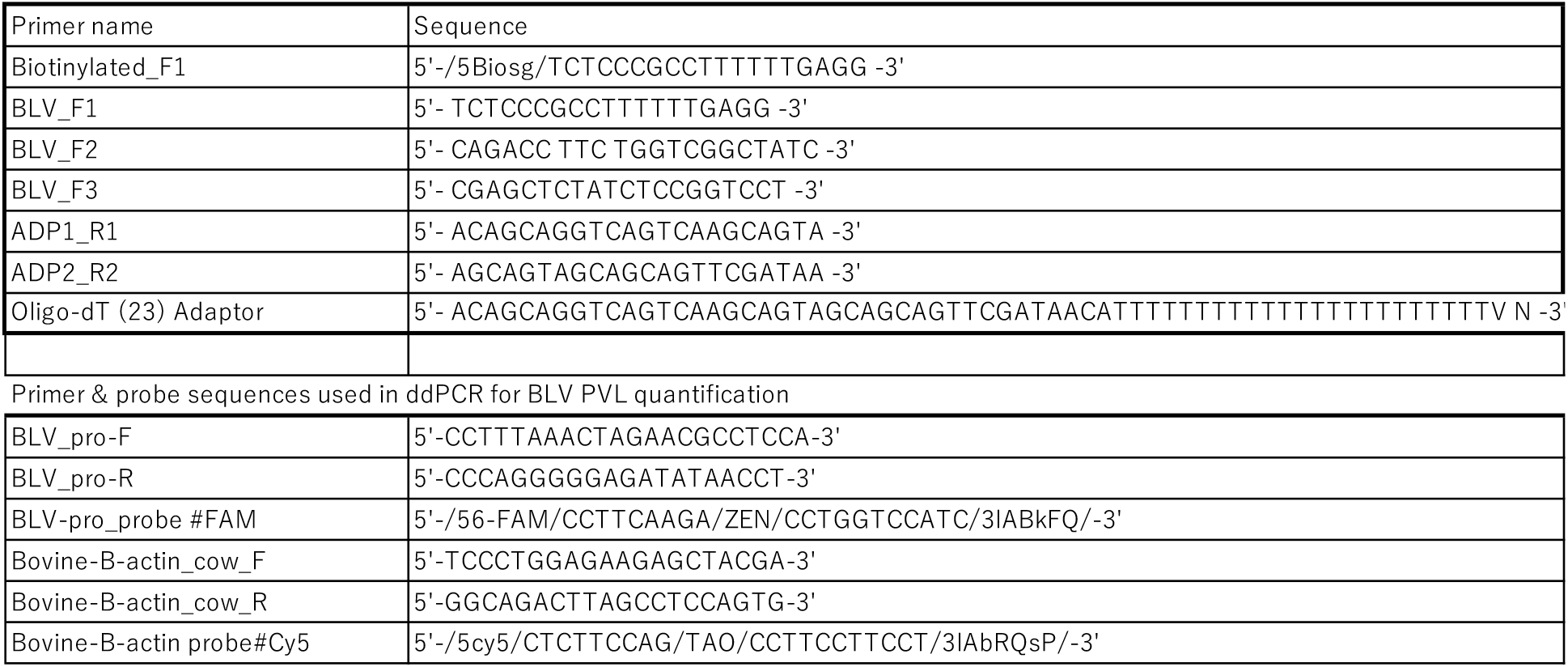
Primer name and their sequences used in BLV-RAIS.

## Supplemental Method

### BLV RAIS Protocol

#### (1) Synthesize ssDNA (Critical step)

Amount of template-500 ng: vortex mix and spin down well before adding to the reaction mix. Imp. Point: Try to adjust DNA conc. below 200 ng/uL. Poor quality DNA templates might interfere with the PCR reaction. We do not recommend using a DNA template with an A260/A230 value lower than 1.5 (optimum is 2 to 2.20).

| Composition | Used(μL) | Final concentration |
| --- | --- | --- |
| 10x PCR Buffer for KOD-plus-Neo | 5 | 1x |
| 2mM dNTPs (each) | 5 | 200μM |
| 25mM MgSO<sub>4</sub> | 3 | 1.5mM |
| 10μM BLV-F1 | 1.5 | 0.3μM |
| KOD-plus-Neo (1U/μL) polymerase | 1 |  |
| Template + H<sub>2</sub>O | 34.5 |  |
| Total | 50 |  |

|  | Temperature | Time | Cycle |
| --- | --- | --- | --- |
| Pre-denaturation | 94°C | 2min | 1 |
| Desaturation | 98°C | 10sec | 25 |
| Extension | 68°C | 43sec |  |

#### (2) Column purification

1. Transfer 50 µL of PCR product to each of the Eppendorf tubes.
2. Add 250 µL (x 5) of Buffer PB and mix well by pipetting.
3. Transfer the total 300 µL to 2 mL QIAquick column.
4. Centrifuge at 12000 RPM for 1 min.
5. Discard the flow-through.
6. Add 730 µL Buffer PE.
7. Centrifuge at 12000 RPM for 1 min.
8. Discard the flow-through.
9. Centrifuge at 12000 RPM for 1 min.
10. Discard the collection tube with flow-through.
11. Place the column on 1.5 mL Lo-Bind Eppendorf tube.
12. Add 30 µL Buffer EB.
13. Centrifuge at 12000 RPM for 1 min.
14. Stored at −20 ℃.

#### (3) PolyA-tailing

| Composition | Used(μL) | Final concentration |
| --- | --- | --- |
| 10x TdT buffer | 1 | 1x |
| 2,5mM CoCL<sub>2</sub> | 1 | 0.25mM |
| 10mM dATP | 0.35 | 0.35mM |
| Terminal Transferase | 0.25 |  |
| ssDNA | 7.4 |  |
| Total | 10 |  |

Incubation: Temp. 37 $^{\circ}$ C. for 30 minutes. => quickly proceed to dsDNA synthesis step

#### (4) Synthesize dsDNA

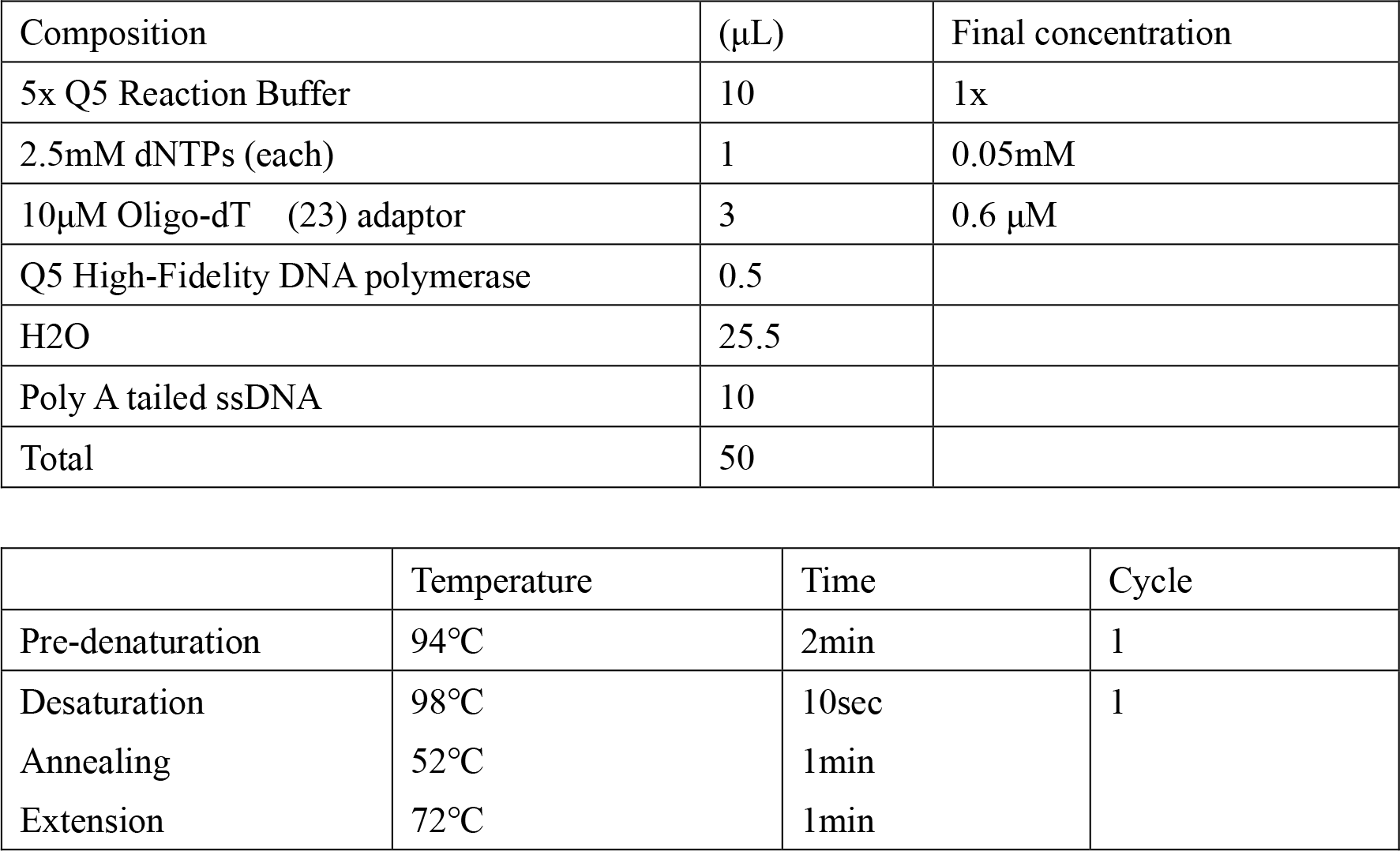

| Composition | (μL) | Final concentration |
| --- | --- | --- |
| 5x Q5 Reaction Buffer | 10 | 1x |
| 2.5mM dNTPs (each) | 1 | 0.05mM |
| 10μM Oligo-dT (23) adaptor | 3 | 0.6 μM |
| Q5 High-Fidelity DNA polymerase | 0.5 |  |
| H<sub>2</sub>O | 25.5 |  |
| Poly A tailed ssDNA | 10 |  |
| Total | 50 |  |

|  | Temperature | Time | Cycle |
| --- | --- | --- | --- |
| Pre-denaturation | 94$^{\circ}$C | 2min | 1 |
| Desaturation | 98$^{\circ}$C | 10sec | 1 |
| Annealing | 52$^{\circ}$C | 1min | |
| Extension | 72$^{\circ}$C | 1min | |

#### (5) First PCR

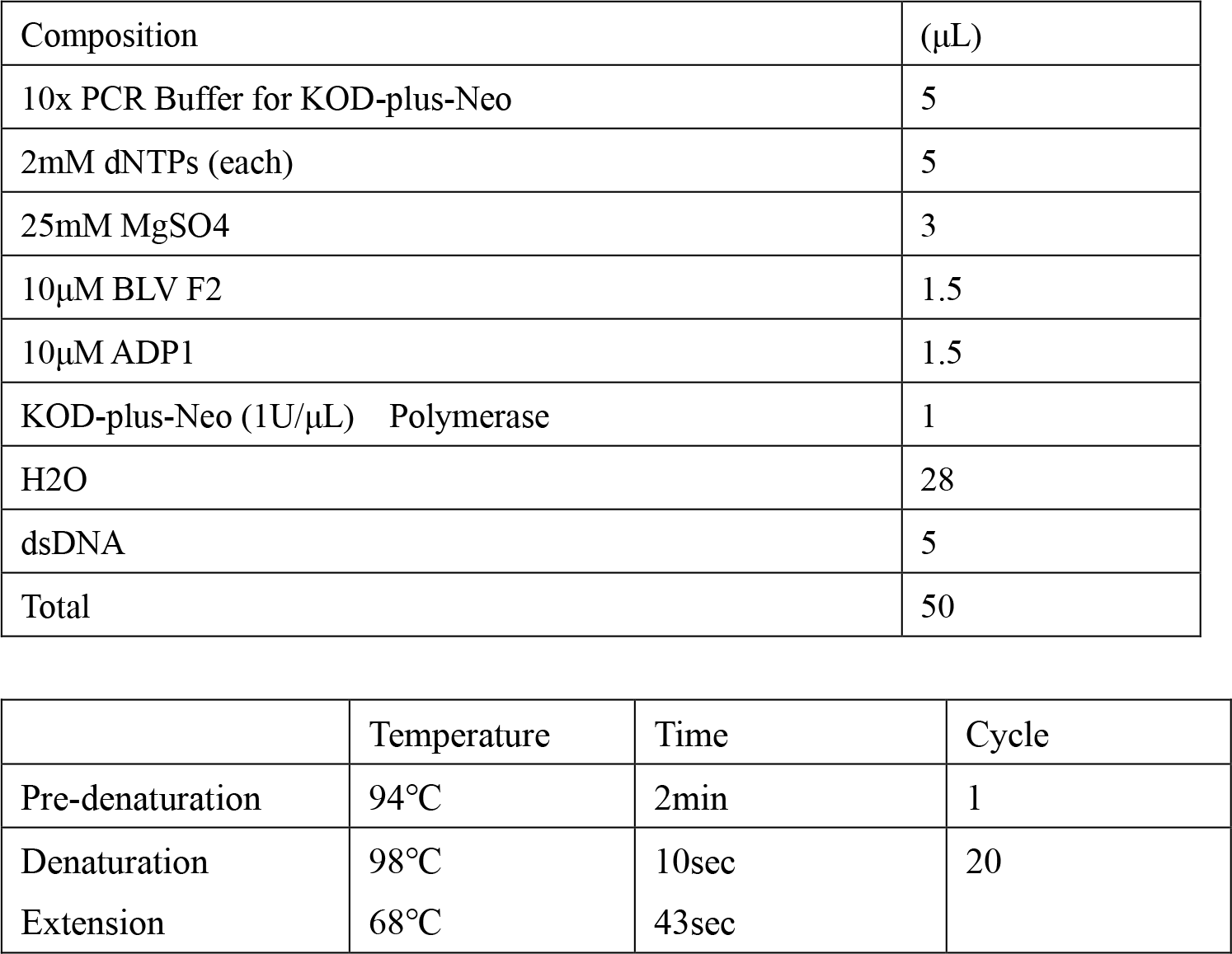

| Composition | (μL) |
| --- | --- |
| 10x PCR Buffer for KOD-plus-Neo | 5 |
| 2mM dNTPs (each) | 5 |
| 25mM MgSO<sub>4</sub> | 3 |
| 10μM BLV F2 | 1.5 |
| 10μM ADP1 | 1.5 |
| KOD-plus-Neo (1U/μL) Polymerase | 1 |
| H<sub>2</sub>O | 28 |
| dsDNA | 5 |
| Total | 50 |

|  | Temperature | Time | Cycle |
| --- | --- | --- | --- |
| Pre-denaturation | 94°C | 2min | 1 |
| Denaturation | 98°C | 10sec | 20 |
| Extension | 68°C | 43sec |  |

#### (6) Second PCR

| Composition | (μL) |
| --- | --- |
| 10x PCR Buffer for KOD-plus-Neo | 5 |
| 2mM dNTPs (each) | 5 |
| 25mM MgSO<sub>4</sub> | 3 |
| 10μM BLV F3 | 1.5 |
| 10μM ADP2 | 1.5 |
| KOD-plus-Neo (1U/μL) Polymerase | 1 |
| H<sub>2</sub>O | 32 |
| First PCR product | 1 (1/100 diluted) |
| Total | 50 |

|  | Temperature | Time | Cycle |
| --- | --- | --- | --- |
| Pre-denaturation | 94°C | 2min | 1 |
| Denaturation | 98°C | 10sec | 20 |
| Annealing | 64°C | 30sec |  |
| Extension | 68°C | 43sec |  |

## Notes

### Competing Interest Statement

The authors have declared no competing interest.

